# Selecting Covariates for Genome-Wide Association Studies

**DOI:** 10.1101/2023.02.07.527425

**Authors:** Erez Dor, Ido Margaliot, Nadav Brandes, Or Zuk, Michal Linial, Nadav Rappoport

## Abstract

The choice of which covariates to include in a Genome-Wide Association Study (GWAS) is important since it affects the ability to detect true association signal of variants, to correct for confounders and avoid false positives, and the running time of the analysis. Commonly used covariates include age, sex, genotyping batches, genotyping array type, as well as an arbitrary number of Principal Components (PCs) used to adjust for population structure. Despite the importance of this issue, there is no consensus or clear guidelines for the right choice of covariates. Therefore, studies typically employ heuristics for their choice with no clear justification. Here, we explore the dependence of the GWAS analysis results on the choice of covariates for a wide range of quantitative and binary human phenotypes. We propose guidelines for covariates choice based on the phenotype’s type (quantitative vs. disease), the heritability, and the disease prevalence, with the goal of maximizing the statistical power to detect true associations and fit accurate polygenic scores while avoiding spurious associations and minimizing computation time. We analyze 36 traits in the UK-Biobank dataset. We show that the genotype batch and assessment center can be safely removed as covariates, thus significantly reducing the GWAS computational burden for these traits.

## 1 Introduction

A central goal in performing Genome-Wide Association Study (GWAS) is to identify statistically significant associations between genetic variations and phenotype and thus point to the possible biological mechanisms underlying the studied phenotype. However, GWAS is often prone to multiple uncontrolled confounders and biases (e.g., selection bias and population structure) [29]. The most common genetic variations tested in GWAS studies are Single Nucleotide Polymorphisms (SNPs). The standard practice in GWAS is to test each SNP independently for association with the trait, which may lead to a high rate of false positives when confounders that are correlated with both the trait and the variant are not included in the model as covariates (i.e. variants that are labeled as statistically associated with the phenotypes but are actually false positives) [2]. The routine GWAS protocol suggests including covariates whose purpose is to control for the indirect effects unrelated to the phenotype of interest and eliminate the influence of confounders. These covariates include technical components (e.g. the genomic center and SNP-chip technology used for collecting data) but also covariates of biological and medical importance, such as the sex and age of the individual, that may directly affect the phenotype.

In recent years, increased GWAS sample sizes and improved statistical methods have lead to an interest in using GWAS results for genomic prediction using Polygenic Scores (PGS). These scores, defined as weighted linear combination of risk alleles, may include SNPs that do not reach genome-wide statistical significance individually, but together can improve prediction accuracy, and were demonstrated as effective for predicting individuals at risk for disease [15]. Thousands of PGSs were already fitted and are available in resources such as [16], with the number of variants included in the score ranging from a few dozens to hundreds of thousands. In similar to the search for genome-wide significant SNPs, the fitting of a PGS may also be susceptible to confounders, and the fitted score will vary depending on the covariates included. A major issue of current interest is the transferability of the scores between different scenarios. In particular, the scores may not transfer easily between human populations [1,24,30], mainly due to differences in allele frequencies, LD-structure, and effect size. Moreover, scores may show reduced accuracy even within a single population where most above differences are negligible [21], including in prediction of within-family variation [27], with changes in covariates such as socioeconomic status, age and sex leading to decreased accuracy, possibly due to Gene-by-Environment interactions. These issues highlight the need to understand the possible confounders affecting the accuracy of the fitted PGSs, and the effect of covariate inclusion on the scores, with the hope that such better understanding will aid in the inclusion of the right covariates when adapting a PGS to a new population or cohort.

Covariates can bias the GWAS results [2], but can also adjust for confounders and prevent spurious associations. For example, population structure has been shown to greatly affect GWAS results [18,13], and including genetic principal components (PCs) as covariates is often used to control for population structure [28,23]. Failure to match cases and controls for the right covariates may also lead to substantial inflation of false positive rate [7,19]. In addition to the effect on the false positive rate, adding covariates may also increase or reduce the statistical power to detect true significant associations [22]. Finally, the addition of covariates to the model comes at a computational price, since multiple regression is performed with the covariates for each SNP repeatedly. Therefore, we may not want to include additional covariates if they do not significantly improve the statistical properties of the analysis.

However, it is often unclear which covariates should be included when performing a GWAS, and what effects will this choice have on the GWAS results [20]. The question of whether under a predetermined setting, a preferred set of covariates should be used is critical to improve the detection power of GWAS while also boosting the accuracy of the findings. To this end, we took an exploratory approach, and performed GWAS for a broad range of traits in the UK-Biobank (UKBB) dataset [5]. For each trait, we performed multiple GWAS with different sets of covariates. We list multiple measures of power and false-positive rates such as the estimated genomic control inflation factor [10] to explore the effect of covariates selection on GWAS. We determine the effect of different covariates for different traits. Specifically, we study how does the heritability of phenotypes influence the effects of different covariates. For binary disease traits (e.g., schizophrenia), we also test the dependence on the disease’s prevalence.

In this study, we propose a criterion for selecting a set of covariates by designing quantitative measures that will enable high discovery power, as well as avoid spurious discoveries. We suggest an optimal set of covariates for different scenarios. Specifically, we will be interested in covariates sets that achieve multiple, possibly competing goals: to minimize the genomic inflation, to maximize the prediction performance, while also minimizing run time.

In this study we propose recommendations for the choice of covariates as a piece of practical advice when dealing with major human quantitative and binary traits in the UK biobank data. By examining many different sets of covariates for dozens of phenotypes in the UK-Biobank dataset, we find that for this dataset, the assessment center and genotyping batch can be excluded from the covariates set without compromising GWAS performance. However, for binary traits, there seems to be some effect of the genotype batch and the assessment center when estimating PGS.

## 2 Methods

### 2.1 GWAS model

Consider a covariates matrix *Z* ∈ ℝ_*n*×*q*_ with *n* individuals (rows) and *q* covariates *Z*_1_,…, *Z*_*q*_ divided into groups *S*_1_, …, *S*_*K*_ (for example, *S*_1_ may contain the Principal Components, *S*_2_ all categorical dummy variables representing assessment center etc.). Consider also the genotypes matrix *X* ∈ ℝ_*n*×*p*_ with SNPs *X*_1_,…, *X*_*p*_, and the phenotypes matrix *Y* ∈ ℝ_*n*×*m*_ with phenotypes *Y*_1_,…, *Y*_*m*_.

We assume a linear model relating a single quantitative phenotype *Y* to known genetic and non-genetic covariates:

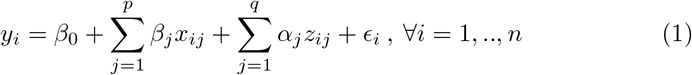

Where ϵ_i_ is an additive noise variable representing environmental effects and other unaccounted-for factors, for individual *i*. In this model, a SNP *j* is termed as true (false) causal if *β*_*j*_ ≠ 0 (*β*_*j*_ ≠ 0). A false causal SNP declared as significant in a GWAS analysis is termed false positive. For disease phenotypes, a similar model is defined using logistic or probit regression.

### 2.2 The Dataset Used

We used the UKBB dataset which includes genotypic and phenotypic data of about 500,000 subjects [5]. We analyzed 36 phenotypes representing a variety of phenotypes with known genetic contribution, for 19 continuous phenotypes and 17 binary disease phenotypes (See Figure 1 and Table 1 for a detailed list). Data from 488,377 samples was used. Samples without the phenotype of interest were filtered out.

**Table 1:**
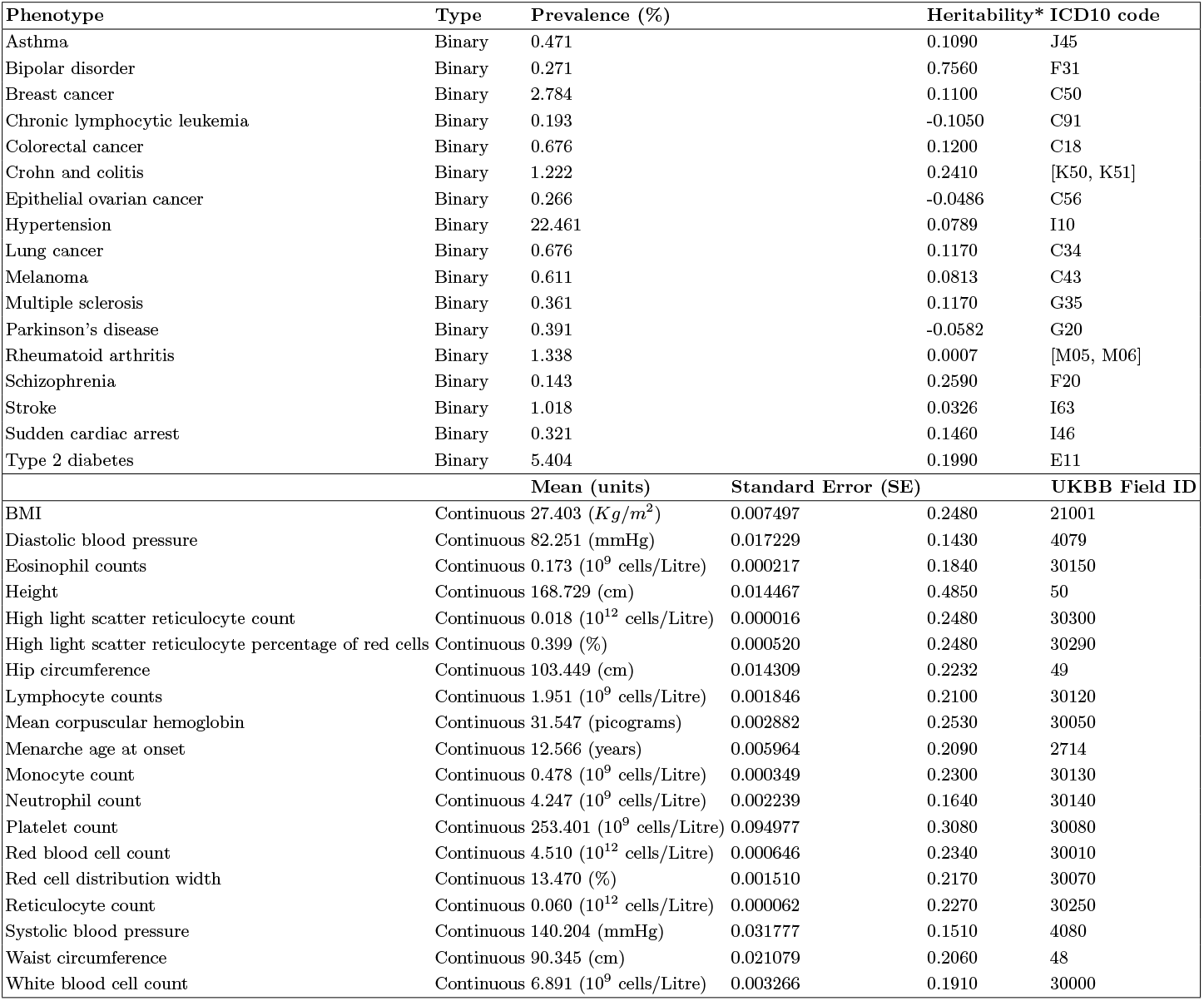
List of phenotypes used in the current study. Prevalence and mean/SE were calculated from the UKBB data in this study. Heritability estimates were taken from Neale lab heritability browser at https://nealelab.github.io/UKBBldsc/index.html.

**Fig. 1:**
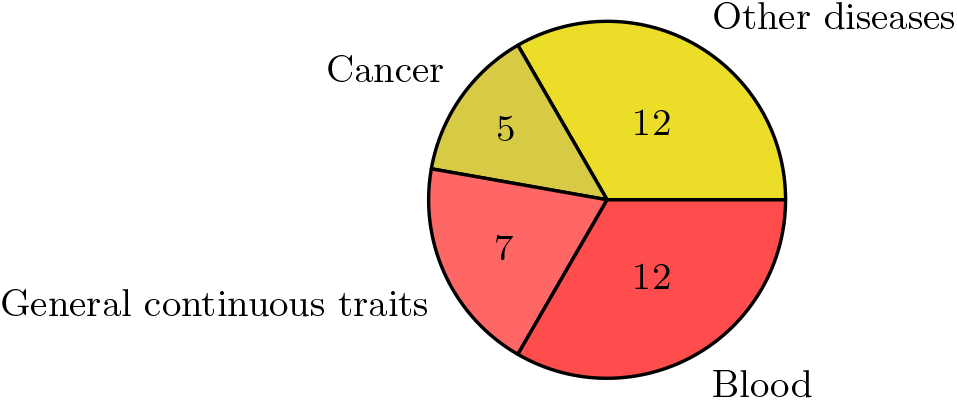
The number of traits in UKBB that were analyzed in this study. In yellow - seventeen binary traits (diseases) and in red - nineteen continuous traits such as blood tests and other physiological measurements. A complete list is shown in Table 1.

For the UKBB dataset, we examined *q* = 168 possible covariates divided into *K* = 6 groups: Genotyping batch, assessment center, sex, age, First 5 PCs, next 35 PCs. We chose to test nine subsets of covariates. Twenty-two assessment centers were represented by 21 binary covariates (dummy variables). Similarly, 106 genotyping batches were represented by 105 binary covariates (see Table 2).

**Table 2:** List of subsets of covariates and number of covariates in each.

| Covariates | Number of covariates |
| --- | --- |
| ∅ | 0 |
| {Age} | 1 |
| {Sex} | 1 |
| {Sex, Age} | 2 |
| {First 5 PCs} | 5 |
| {Sex, Age, 40 PCs} | 42 |
| {Sex, Age, Assessment Center, 40 PCs} | 63 |
| {Sex, Age, Batch, 40 PCs} | 147 |
| {Sex, Age, Batch, Assessment Center, 40 PCs} | 168 |

### GWAS execution

GWAS was performed using Plink2 [6][26] using `“--glm” command. Covariates were standardized using the “--covar-variance-standardize”` flag.

### 2.3 Evaluation metrics

As ground truth for GWAS is not available, we considered multiple evaluation metrics for assessing the quality and optimality of covariates’ selection.

### Polygenic scores accuracy

For each pair of a phenotype and a covariates subset, we trained a PGS model using the GWAS’s summary statistics (additive effect size *β*_*j*_ and significance level (*Pval*_*j*_) of every SNP *j*) with this set of covariates. PGS was computed using PRSice-2 software [12]. The score is estimated by removing SNPs in linkage disequilibrium (LD) and by thresholding the p-values, where an optimal p-value threshold is chosen to optimize the prediction accuracy of the resulting PGS.

We used as an evaluation metric the percent of phenotypic variance explained (*R*^2^) by the PGS [11], a metric indicative of the prediction quality of the PGS model trained based on the GWAS results with a particular covariates set [3]. *R*^2^ was computed by running PRSice-2 using all samples. The same subset of covariates that was used to estimate the effect sizes was also used for estimating *R*^2^ of the PGS. In total, we trained 324 PGS models (36 phenotypes } 9 covariates sets). The full *R*^2^ represents the variance explained by both the PGS and the covariates.

#### Genomic control inflation factor

To control for inflation of discoveries in GWAS, [10] introduced the λ inflation factor statistic and proposed a method called ’genomic control’, which utilizes this statistic to correct for false positive signals. We used the λ inflation factor as an evaluation metric that measures the discrepancy between the empirical p-value distribution and the null *Uniform*(0, 1) distribution. This statistic is defined as the scaled median of the individual SNPs’ χ^2^ test statistics, with λ = 1 indicating a complete agreement and a higher value indicating a large number of significant associations that can be due to confounders and/or a true polygenic signal. Genomic λ inflation factor was computed using PLINK-2 [6].

#### Linkage Disequilibrium Score Regression’s Intercept

We run LD Score Regression (LDSC) [4] to discriminate between confounders and a true polygenic score for each pair of phenotype and covariates-subset. The measure of interest was the intercept of LDSC-regression (LDSCI). LDSCI provides an estimate of the confounder effect [4] based on a simple yet powerful idea. Since SNPs are correlated due to linkage-disequilibrium, the signal observed in GWAS for a single SNP can be a proxy for the signal of other neighboring SNPs. The LD-score of a SNP measures the cumulative correlations between this SNP and neighboring SNPs. For a true polygenic signal that is spread across many SNPs, we expect, on average more causal SNPs nearby a SNP with a higher LD-score, hence a linear relationship between the χ^2^ association statistic of a SNP and its LD-score. In contrast, we expect confounders such as population structure to affect SNPs more uniformly and independently of their LD-score. Therefore, the true polygenic signal is correlated with the LD-score of a SNP and is reflected in the slope of the regression line between the LD-score and the χ^2^ association statistic. In contrast, the intercept of this regression analysis reflects the inflation due to confounders. This metric measures the level of spurious associations in a GWAS, with *LDSCI* = 1 indicative of no confounding, and values above 1 indicate confounding (see [17] for additional details). In contrast to the genomic control λ, this metric is not inflated by a true polygenic signal that is correlated with the LD-level of individual SNPs. For computing LDSC we removed strand-ambiguous SNPs, and used the European population from the 1,000 Genomes project as a reference panel [8] and for computing the LD scores.

## 3 Results

A diverse set of 36 phenotypes was selected for this study covering a variety of traits (Figure 1): 17 disease traits of different prevalence (five cancer diagnoses and twelve diagnoses of other diseases) and 19 continuous phenotypes (twelve blood test results and seven other continuous physiological measurements). The number of cases for the different binary traits ranges from 62,000 for hypertension to only 127 for systemic sclerosis. More details about the phenotypes are shown in Table 1. For each phenotype, we executed multiple GWAS with nine different sets of confounders, as shown in Table 2.

### 3.1 Genome-wide Significant SNPs

A possible measure of GWAS success is the number of genome-wide significant SNPs. We first recorded this number for each combination of covariates and phenotypes at *α* = 5 × 10^−8^ [9]. The number of genome-wide significant SNPs for each trait-covariates combination is shown in Figure 2. The color of each cell represents the percentage of significant SNPs compared with the base value on the left-most column, which is the number of significant SNPs without any covariates.

**Fig. 2:**
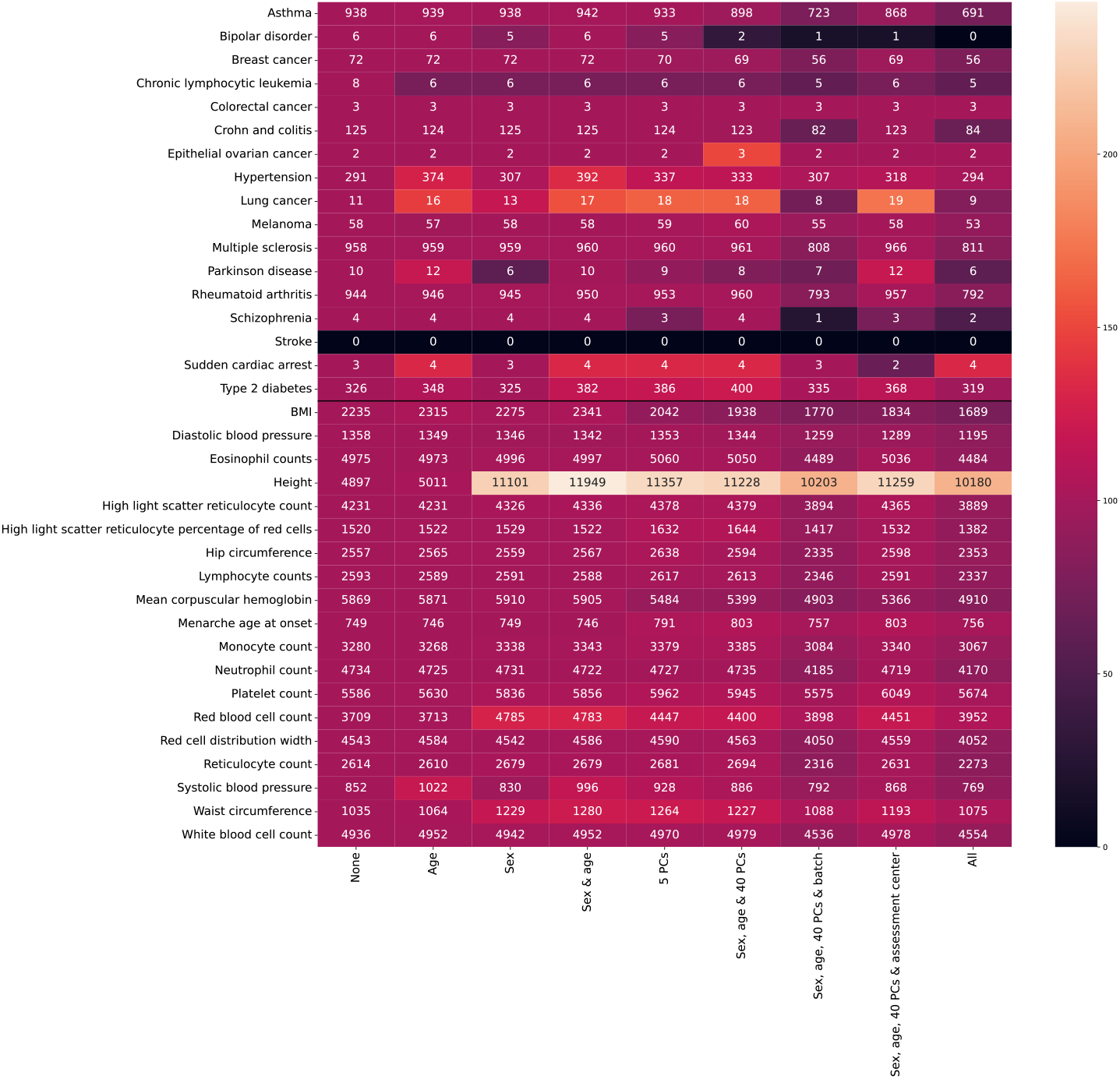
The number of genome-wide significant SNPs for each phenotype-covariates combination. The color of each cell represents the percentage of significant SNPs compared to the base value on the leftmost column, which is the number of significant SNPs without any covariates. The top rows are binary traits and the bottom rows are continuous.

### 3.2 Genomic Inflation factor

We computed λ inflation factor across all (phenotype, covariates-subset) pairs. Usually, adding covariates to the association test decreases this inflation factor. As a rule of thumb, λ < 1.1 is considered acceptable [31]. Therefore, we hypothesized that the subset of covariates that minimizes inflation is more adequate as it controls for the correct confounders of the population. However, adding more covariates may lead to overcorrection and missing true SNPs while lowering the λ value. The λ inflation factors for each run are summarized in Table 3.

**Table 3:** λ inflation factor for each set of covariates for each phenotype, using the Plink2 software. Upper part contains binary phenotypes and lower part contain continuous phenotypes. Minimal value in each row is in bold face. In each row, the values within 105% from the minimal values are highlighted in magenta. AC-Assessment center; Batch-Genotype measurement batch.

|  | None | Age | Sex | Sex & Age | 5Pcs | W/O Batch & AC | W/O AC | W/O Batch | All |
| --- | --- | --- | --- | --- | --- | --- | --- | --- | --- |
| Asthma | 1.162 | 1.162 | 1.162 | 1.165 | 1.159 | 1.143 | **1.14** | 1.143 | **1.14** |
| Bipolar Disorder | **1.02** | **1.02** | **1.02** | **1.02** | **1.02** | **1.02** | 1.023 | **1.02** | **1.02** |
| Breast Cancer | 1.083 | **1.08** | 1.083 | **1.08** | **1.08** | **1.08** | 1.083 | **1.08** | **1.08** |
| Chronic Lymphocytic Leukemia | **1.023** | **1.023** | **1.023** | **1.023** | 1.026 | **1.023** | **1.023** | **1.023** | **1.023** |
| Colorectal Cancer | 1.047 | 1.047 | 1.047 | 1.047 | **1.044** | **1.041** | 1.044 | **1.041** | 1.044 |
| Crohn And Colitis | 1.047 | 1.047 | 1.047 | 1.047 | 1.044 | 1.044 | **1.038** | 1.044 | **1.038** |
| Epithelial Ovarian Cancer | **1.005** | **1.002** | 1.005 | **1.002** | **1.002** | **1.002** | 1.005 | **1.002** | 1.005 |
| Hypertension | 1.398 | 1.418 | 1.389 | 1.418 | 1.396 | 1.355 | 1.351 | 1.355 | **1.337** |
| Lung Cancer | 1.029 | 1.029 | 1.029 | 1.029 | **1.02** | **1.02** | 1.026 | **1.02** | 1.023 |
| Melanoma | 1.032 | 1.032 | **1.029** | 1.032 | 1.032 | 1.032 | 1.032 | 1.032 | 1.032 |
| Multiple Sclerosis | **1.026** | 1.029 | 1.029 | 1.029 | **1.026** | **1.026** | 1.032 | **1.026** | 1.032 |
| Parkinson Disease | 1.029 | 1.029 | 1.032 | **1.023** | 1.026 | 1.032 | 1.029 | 1.032 | 1.029 |
| Rheumatoid Arthritis | **1.056** | **1.056** | **1.056** | 1.056 | **1.053** | **1.053** | **1.053** | **1.053** | **1.053** |
| Schizophrenia | 1.008 | 1.008 | 1.011 | 1.011 | **1.002** | 1.008 | 1.011 | 1.008 | 1.011 |
| Stroke | **1.014** | **1.011** | **1.011** | 1.014 | **1.011** | **1.011** | **1.011** | **1.011** | **1.011** |
| Sudden Cardiac Arrest | **1.005** | **1.005** | **1.005** | **1.005** | **1.005** | 1.008 | 1.008 | 1.008 | **1.005** |
| Type 2 Diabetes | 1.22 | 1.22 | 1.217 | 1.217 | 1.217 | 1.21 | **1.207** | 1.21 | **1.207** |
| Bmi | 1.893 | 1.901 | 1.897 | 1.91 | 1.832 | 1.816 | 1.797 | 1.816 | **1.785** |
| Diastolic Blood Pressure | 1.46 | 1.464 | 1.46 | 1.457 | 1.457 | 1.446 | 1.446 | 1.446 | **1.432** |
| Eosinophil Counts | 1.403 | 1.403 | 1.407 | 1.407 | **1.375** | **1.372** | **1.369** | 1.372 | 1.372 |
| Height | **1.988** | 2.034 | 2.67 | 2.842 | 2.26 | 2.172 | 2.159 | 2.172 | 2.142 |
| High Light Scatter Reticulocyte Count | 1.533 | 1.536 | 1.544 | 1.547 | 1.507 | 1.5 | 1.496 | 1.5 | **1.482** |
| High Light Scatter Reticulocyte Percentage Of Red Cells | 1.256 | 1.256 | 1.256 | 1.256 | 1.23 | 1.23 | 1.227 | 1.23 | **1.22** |
| Hip Circumference | 1.741 | 1.741 | 1.741 | 1.741 | 1.737 | 1.73 | 1.722 | 1.73 | **1.71** |
| Lymphocyte Counts | 1.24 | 1.243 | 1.24 | 1.24 | 1.227 | **1.22** | **1.214** | 1.22 | **1.214** |
| Mean Corpuscular Hemoglobin | 1.562 | 1.577 | 1.577 | 1.588 | 1.386 | **1.369** | **1.369** | **1.369** | 1.372 |
| Menarche Age At Onset | 1.425 | 1.421 | 1.425 | 1.421 | 1.414 | **1.403** | **1.403** | **1.403** | **1.403** |
| Monocyte Count | 1.31 | 1.317 | 1.317 | 1.32 | 1.307 | 1.307 | **1.3** | 1.307 | **1.3** |
| Neutrophil Count | 1.507 | 1.507 | 1.507 | 1.507 | 1.496 | 1.482 | 1.475 | 1.482 | **1.471** |
| Platelet Count | 1.668 | 1.656 | 1.675 | 1.672 | 1.592 | 1.581 | **1.573** | 1.581 | **1.573** |
| Red Blood Cell Count | 1.649 | 1.649 | 1.73 | 1.733 | 1.581 | **1.573** | **1.566** | 1.573 | 1.57 |
| Red Cell Distribution Width | 1.3 | 1.303 | 1.303 | 1.303 | 1.3 | 1.303 | 1.297 | 1.303 | **1.293** |
| Reticulocyte Count | 1.29 | 1.29 | 1.293 | 1.293 | 1.29 | 1.283 | **1.28** | 1.283 | **1.276** |
| Systolic Blood Pressure | **1.449** | 1.482 | **1.449** | 1.489 | 1.485 | 1.471 | 1.467 | 1.471 | 1.457 |
| Waist Circumference | **1.607** | **1.615** | 1.718 | 1.733 | 1.733 | 1.714 | 1.702 | 1.714 | 1.691 |
| White Blood Cell Count | 1.507 | 1.507 | 1.504 | 1.507 | 1.489 | 1.478 | **1.471** | 1.478 | **1.464** |

### 3.3 Polygenic Score Accuracy

We evaluated the predictive power of the different GWAS results as used in Polygenic risk scores. For every phenotype and covariates-subset pair, we add the same set of covariates when computing the PGS’s *R*^2^. We found that for most binary phenotypes (12 out of 17), using all covariates maximizes the *R*^2^. The second best covariates subset was ’without genotype batch’. The subset of covariates ’without genotype batch and assessment center’ achieved *R*^2^ within between 98% and 100% of the maximal *R*^2^ for all phenotypes except for Colorectal cancer for which it achieved 62% of the maximal *R*^2^ of 0.58 which was found only using all covariates (Table 4).

**Table 4:**
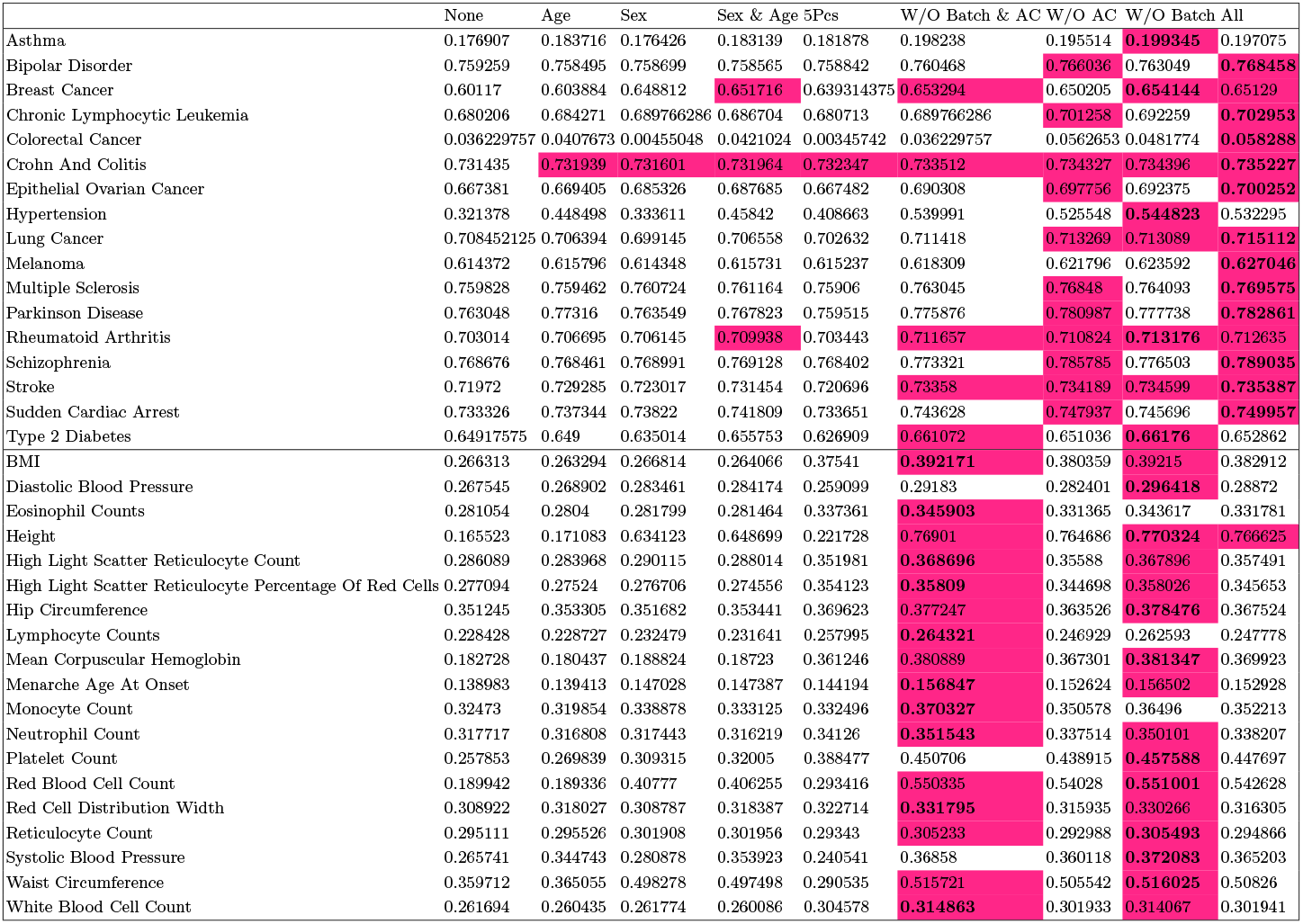
Percent variance explained (R^2^) of PGS for each set of covariates for each phenotype. The R^2^ indicates the predictive power of each set of covariates. The upper part contains binary phenotypes and lower part contain continuous phenotypes. Highest value in each phenotype (row) is marked in bold. In each row, the values within 99.5% from the maximal values are highlighted. AC-Assessment center; Batch-Genotype measurement batch

For quantitative phenotypes, removing either the genotype batch or the assessment center improved the *R*^2^ compared to using all covariates, and one of the subsets ’without genotype batch and assessment center’ and ’without genotype batch’ yielded the maximal *R*^2^ for all 19 phenotypes.

### 3.4 LDSCI

We next calculated the LD-Score regression’s intercept (LDSCI) using LDSC v1.0.1 [4]. The LDSCI values for each run are summarized in Table 5. Overall, the results are similar to the result of the genomic control metric. For binary traits the effect of covariates on the LDSCI metric is minimal for most traits except Asthma, Hypertension, and Lung cancer, where the exclusion of Principal Components decreases the LDSCI. For continuous traits, the inclusion of Principal Components seems more critical and may decrease substantially the LDSCI metric.

**Table 5:** Intercept of LDSC for each set of covariates for each phenotype, using the LDSC software. Upper part contains binary phenotypes and lower part contains continuous phenotypes. Minimal value in each row is in bold face. In each row, the values within 105% from the minimal values are highlighted in magenta. AC-Assessment center; Batch-Genotype measurement batch.

|  | None | Age | Sex | Sex & Age | 5Pcs | W/O Batch & AC | W/O AC | W/O Batch | All |
| --- | --- | --- | --- | --- | --- | --- | --- | --- | --- |
| Asthma | 1.017 | 1.017 | 1.018 | 1.018 | 1.016 | 1.005 | 1.004 | 1.004 | **1.003** |
| Bipolar Disorder | **0.99** | **0.99** | **0.99** | **0.99** | **0.99** | **0.99** | 0.994 | 0.991 | 0.994 |
| Breast Cancer | 1.014 | 1.013 | 1.014 | 1.013 | 1.013 | **1.012** | **1.012** | **1.012** | 1.013 |
| Chronic Lymphocytic Leukemia | **0.981** | 0.982 | 0.982 | 0.982 | 0.982 | 0.982 | 0.985 | 0.982 | 0.984 |
| Colorectal Cancer | 0.999 | 1.0 | 1.0 | 1.0 | **0.997** | 0.998 | 0.998 | **0.997** | 0.998 |
| Crohn And Colitis | 1.028 | 1.028 | 1.028 | 1.028 | 1.028 | **1.027** | 1.029 | **1.027** | 1.029 |
| Epithelial Ovarian Cancer | 0.986 | **0.985** | 0.986 | **0.985** | 0.986 | 0.986 | 0.988 | 0.986 | 0.988 |
| Hypertension | 1.091 | 1.089 | 1.087 | 1.088 | 1.071 | 1.046 | 1.048 | **1.043** | 1.045 |
| Lung Cancer | 0.999 | 1.002 | 0.999 | 1.002 | 0.994 | **0.993** | **0.993** | 0.994 | 0.994 |
| Melanoma | 1.013 | 1.013 | 1.013 | 1.013 | 1.011 | 1.011 | **1.007** | 1.01 | **1.007** |
| Multiple Sclerosis | 1.028 | 1.028 | 1.028 | 1.028 | 1.028 | 1.027 | **1.026** | 1.027 | **1.026** |
| Parkinson Disease | **0.993** | **0.993** | **0.993** | 0.994 | 0.995 | 0.994 | 0.998 | 0.995 | 0.999 |
| Rheumatoid Arthritis | 1.019 | 1.019 | 1.02 | 1.02 | **1.018** | **1.018** | 1.019 | **1.018** | **1.018** |
| Schizophrenia | 0.991 | 0.991 | 0.992 | 0.991 | 0.99 | 0.99 | **0.989** | 0.991 | 0.99 |
| Stroke | 1.003 | 1.002 | 1.003 | 1.003 | 1.002 | **1.001** | 1.004 | 1.002 | 1.005 |
| Sudden Cardiac Arrest | **1.0** | **1.001** | **0.999** | **1.0** | **1.0** | **1.0** | 1.001 | 1.001 | 1.001 |
| Type 2 Diabetes | 1.033 | 1.033 | 1.033 | 1.034 | 1.034 | 1.031 | 1.029 | 1.029 | **1.027** |
| Bmi | 1.144 | 1.15 | 1.149 | 1.154 | 1.1 | **1.083** | 1.084 | **1.078** | 1.079 |
| Diastolic Blood Pressure | 1.063 | 1.064 | 1.066 | 1.066 | 1.065 | 1.056 | **1.055** | 1.053 | **1.05** |
| Eosinophil Counts | 1.134 | 1.135 | 1.136 | 1.137 | 1.106 | **1.105** | **1.1** | 1.105 | 1.101 |
| Height | 1.392 | 1.428 | 1.664 | 1.757 | 1.367 | 1.296 | 1.294 | **1.291** | **1.287** |
| High Light Scatter Reticulocyte Count | 1.096 | 1.099 | 1.104 | 1.106 | 1.076 | 1.073 | **1.067** | 1.072 | **1.066** |
| High Light Scatter Reticulocyte Percentage Of Red Cells | 1.048 | 1.049 | 1.048 | 1.05 | 1.028 | 1.025 | **1.02** | 1.024 | 1.022 |
| Hip Circumference | 1.105 | 1.105 | 1.105 | 1.105 | 1.099 | 1.091 | 1.089 | 1.088 | **1.087** |
| Lymphocyte Counts | 1.06 | 1.06 | 1.058 | 1.059 | **1.051** | 1.048 | 1.047 | 1.047 | **1.046** |
| Mean Corpuscular Hemoglobin | 1.22 | 1.229 | 1.23 | 1.24 | 1.093 | **1.077** | 1.081 | 1.079 | 1.082 |
| Menarche Age At Onset | 1.037 | 1.037 | 1.037 | 1.037 | 1.028 | **1.017** | 1.019 | 1.02 | 1.018 |
| Monocyte Count | **1.08** | 1.085 | 1.081 | 1.087 | **1.077** | 1.076 | **1.075** | 1.077 | 1.076 |
| Neutrophil Count | 1.092 | 1.094 | 1.093 | 1.095 | 1.086 | 1.077 | **1.072** | 1.075 | **1.072** |
| Platelet Count | 1.216 | 1.206 | 1.214 | 1.208 | 1.138 | 1.13 | **1.126** | 1.132 | 1.129 |
| Red Blood Cell Count | 1.252 | 1.253 | 1.267 | 1.27 | 1.157 | 1.15 | **1.145** | 1.151 | 1.146 |
| Red Cell Distribution Width | 1.036 | 1.035 | **1.034** | 1.035 | **1.034** | 1.033 | 1.03 | 1.031 | **1.029** |
| Reticulocyte Count | **1.023** | **1.023** | 1.025 | 1.025 | 1.025 | 1.022 | 1.022 | **1.018** | **1.018** |
| Systolic Blood Pressure | 1.089 | 1.096 | 1.089 | 1.096 | 1.093 | 1.085 | 1.086 | **1.079** | 1.08 |
| Waist Circumference | **1.061** | **1.066** | 1.083 | 1.091 | 1.087 | 1.077 | 1.078 | 1.071 | 1.071 |
| White Blood Cell Count | 1.116 | 1.118 | 1.116 | 1.119 | 1.1 | **1.091** | **1.088** | 1.092 | **1.088** |

### 3.5 Comparison across covariate subsets

Figure 3 shows the distribution of the evaluation metrics for each covariate subset across the phenotypes, thus enabling a high-level view of the effects of covariate subsets on the evaluation metrics across phenotypes. For example, for continuous phenotypes, LDSCI decreases when adding more covariates and saturates when 40 PCs are included. A similar observation can be seen when considering PGS’s *R*^2^.

**Fig. 3:**
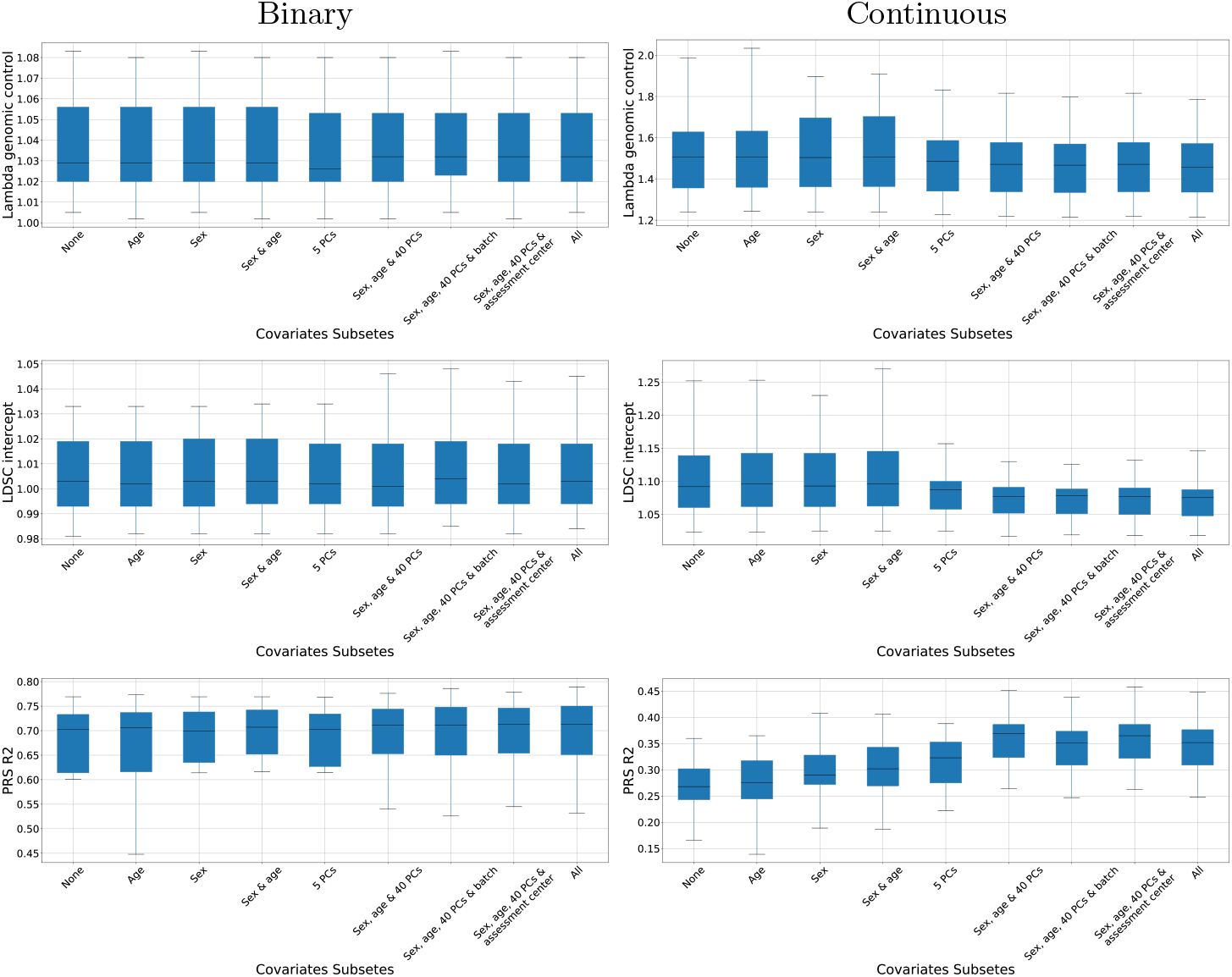
Evaluation metrics summary phenotypes. Top row: λ inflation factor; Middle row: LDSCI, Bottom row: PGS’s *R*^2^. Left columns: binary traits, Right columns: continuous traits. The box plots are per subset of covariates across the different phenotypes.

## 4 Conclusions

In this study, we conducted an empirical evaluation of the set of covariates included in a GWAS study on various metrics representing the GWAS results.

Based on our exploratory analysis, we set to determine a practical recommendation for the choice of covariates to include in the GWAS analysis. The goal is to minimize the *LDSCI*, λ-control, and running time while maximizing the number of genome-wide significant discoveries and the PGS *R*^2^. Based on these criteria, we recommend that PGS estimations include age, sex, and all 40 principal components as covariates for both binary and quantitative traits. It balanced between getting a λ inflation factor close to the minimum, and minimized LDSCI, while PGS *R*^2^ is maximized (Figure 3). For binary traits, the effect of the genotype batch and assessment center seems to be more critical in terms of the PGS’s *R*^2^, and we recommend including all covariates of the analysis if this is the goal of the study. That makes sense if genotyping batch and assessment center have no correlation with the genotype, but are correlated with the trait.

While our recommendations are applicable to the UKBB dataset, our empirical approach can be utilized to suggest the set of covariates for other GWAS studies in other cohorts as well, with differences in population structure, sample size, case-control balances, etc. Specifically, our approach suggests balancing the running time and the statistical properties of the results. The running time is quadratic in the number of covariates and could increase substantially, especially when including multiple PCs and dummy variables with many categories such as batches and assessment centers - hence it is desirable to limit the number of covariates if there is no evidence for significant changes in the GWAS results.

A recent study [25] explored the choice of covariates for GWAS in the UKBB dataset. However, this study focused only on Principal Components and their effect on population structure, concluding that 16-18 PCs should be taken for the UKBB population. The authors did not consider the effect of phenotype on the choice of covariates. Our analysis extends these findings, showing that multiple PCs are indeed required, but also exploring the effect on the phenotype on the choice of covariates, the differences between binary traits and quantitative traits, and the inclusion of additional covariates such as batch and assessment center.

## 5 Acknowledgements

This research has been conducted using the UK Biobank Resource under Application Number 56774.

