## Supplementary for "Selecting Covariates for Genome-Wide Association Studies"

### Appendix A

#### A.1 Comparison between LDSCI and the number of significant SNPs

Next, we compared the LDSCI and the number of genome-wide significant SNPs. Our goal was to test whether adding more covariates reduces the bias confounding (decreasing the LDSC intercept) by controlling for false positive variants. Each panel in Figure A1 represents a phenotype. The X-axis represents the LDSC intercept, and the Y-axis is the number of significant SNPs compared to the number of significant SNPs for the 'None' covariates subset (in percentages). In each panel, the number of SNPs of the 'None' covariates subset is represented as 100% and the other covariates subsets' percent are related to it. Each percent number of different covariates subset from "None" can be $\leq 100\%$ or $> 100\%$. In most phenotypes, including all the covariates gives sub-optimal results - much fewer variants than the optimum, without improving the intercept significantly and, in some cases, even increasing it.

#### A.2 Running Time

We examined the runtime of performing GWAS using each set of covariates (Figure A2). In general, the effect of covariates on the running times for continuous and binary phenotypes was similar. For both trait types, adding more covariates increases the runtime by up to roughly two orders of magnitude compared to the running time without covariates. Tough as expected, the absolute running time of binary traits was around $3 - 5$-times slower.

#### A.3 Comparison to previous work

To test the robustness of our findings to preprocessing choices, we compared our results to the analysis performed by the Neal lab for the UKBB datasets, available at http://www.nealelab.is/uk-biobank, with a focus on heritability estimates [14]. The set of covariates chosen in Neal's analysis is slightly different from our covariates and contains age, age$^2$, sex, age$\times$sex, age$^2\times$sex, and the first 20 PCs. We choose sex, age, and first 40 PCs as covariates for results analysis comparison since this is the most similar to Neale's chosen covariates set. We compared the $\lambda$ inflation factor and LDSCI metrics.

Overall, our results are similar to previous findings by Neal's lab, with a slight decrease in $\lambda$ inflation factor and $LDSCI$ in our analysis, as shown in Figure A3. With this global agreement, we can interpret the heritability estimates from Neal's analysis (see Table 1, together with our own computed metrics.

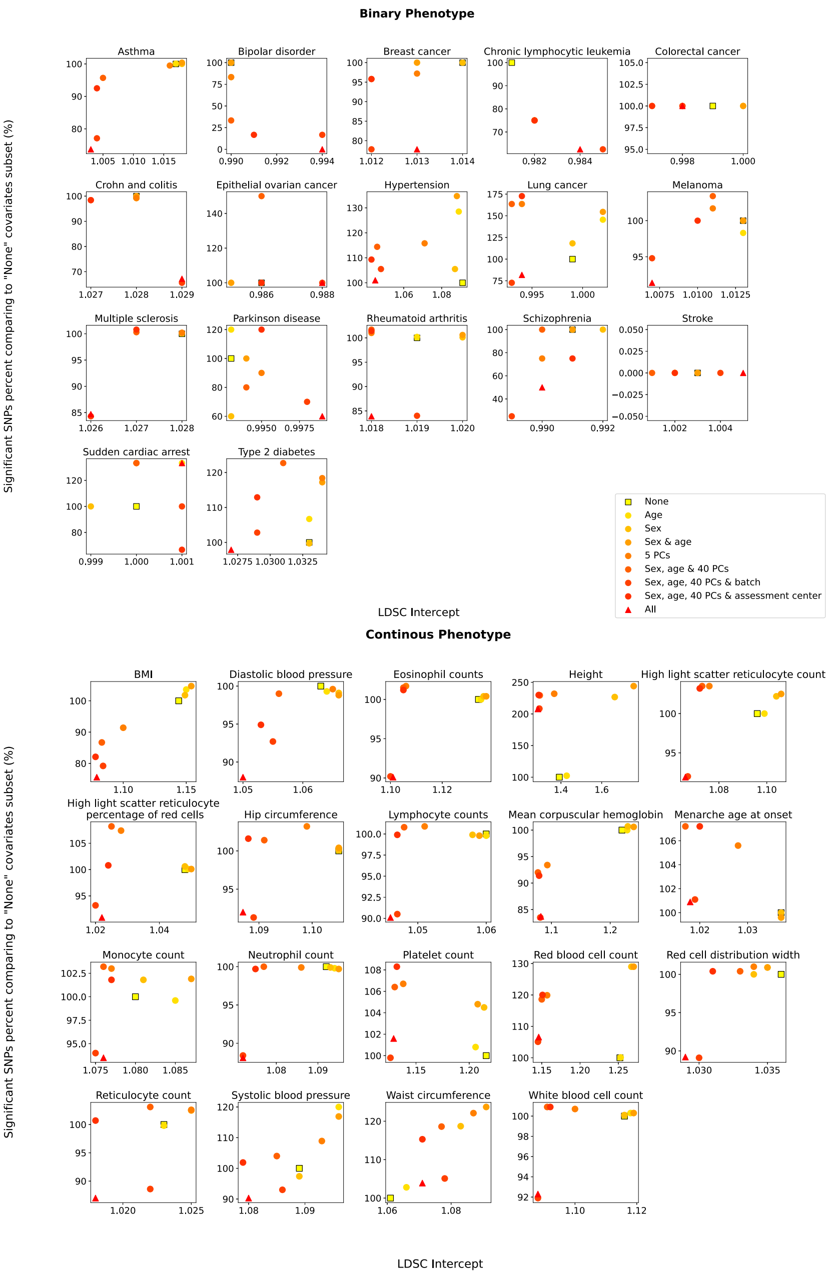


Fig. A1: Significant SNPs percent compared to LDSC intercept. Each panel is a phenotype. The X-axis represents the LDSC intercept, and the Y-axis the percentage of significant SNPs compared to analysis when no covariates were considered. Top: binary phenotypes; Bottom: continuous phenotypes.

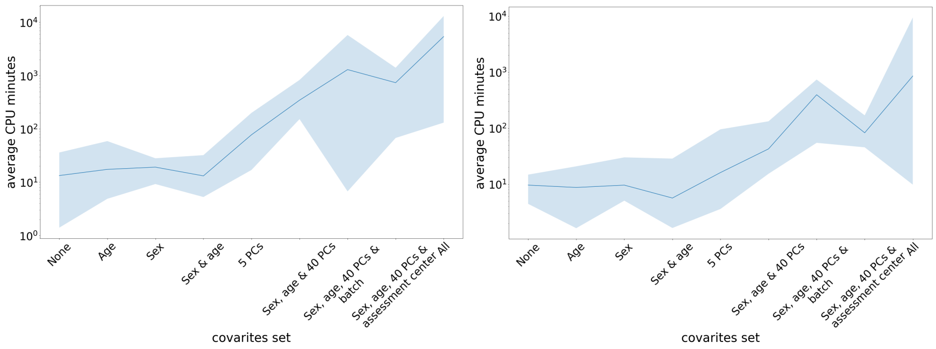


Fig. A2: Left: Run time for binary traits; Right: Run time for continuous traits. To illustrate, we plot the average and standard deviation of performing GWAS for all SNPs of chromosome 1 over 324 runs for each set of covariates (9) and over the different traits (36)

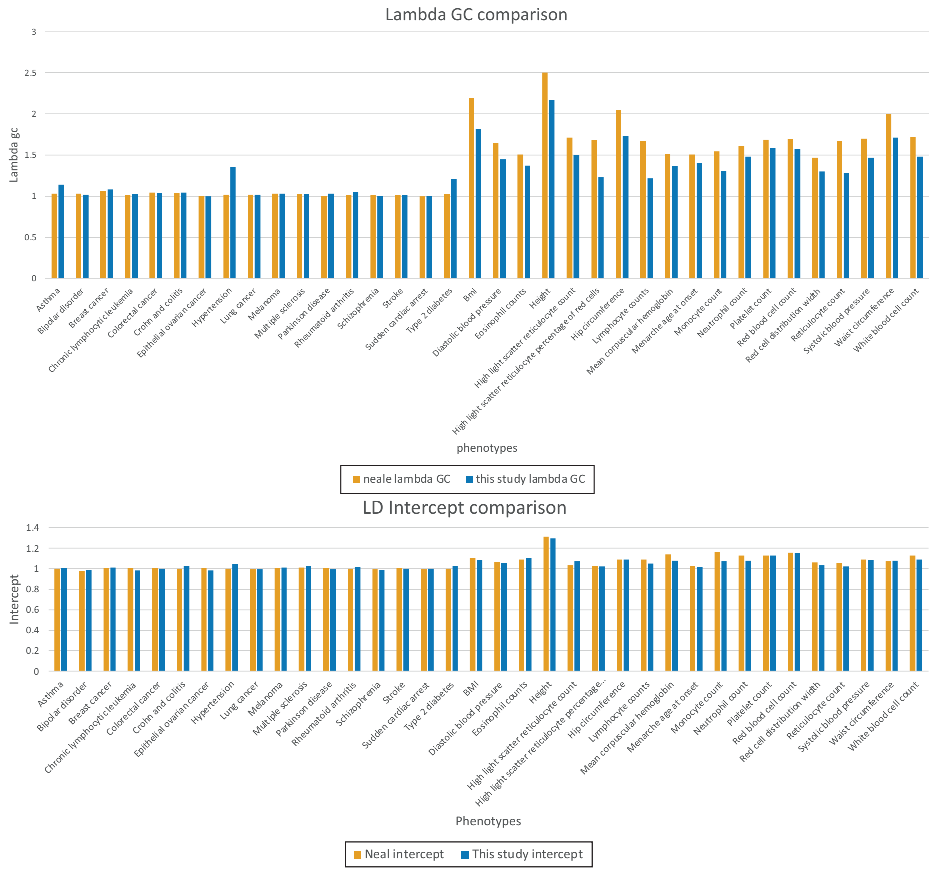


Fig. A3: Differences in $\lambda$ inflation factor (Top) and $LDSCI$ (Bottom) between our analysis and Neal's lab result. The results of both analyses largely agree, with a slightly higher value of $\lambda$ for Neal's lab analysis for continuous traits.
